# PCSK9 Exhibits Novel Nuclear Localization in LSEC and Its Targeting with Bioinspired Nanoparticles Reduces Colorectal Liver Metastasis

**DOI:** 10.64898/2026.05.26.727886

**Authors:** Ander Martin, Maitane Duarte García-Escudero, Helena García García, Iratxe Bañares, Naroa Fontal, Jokin Eguia, Patricia García-Gallastegui, Asier Benito-Vicente, Francisco José Sáez, Olatz Crende, Alejandro Sánchez-Barreiro, Joana Marquez, Abdel-Majid Khatib, Iker Badiola

## Abstract

**Background & Aims:** Colorectal cancer liver metastasis is the leading cause of mortality in affected patients, with liver sinusoidal endothelial cells playing a pivotal role in metastatic niche formation. Proprotein convertase subtilisin/kexin type 9 has emerged as a regulator of tumor biology, but its function in the hepatic microenvironment remains poorly defined. This study aimed to characterize the role and subcellular localization of PCSK9 in liver sinusoidal endothelial cells and to evaluate the therapeutic potential of its endothelial-specific inhibition in colorectal liver metastasis.

**Methods:** *In vitro* studies were performed using human and murine liver sinusoidal endothelial cells stimulated with conditioned media from metastatic colorectal cancer cells and cancer stem cells. Subcellular localization was assessed by immunofluorescence, immunogold electron microscopy, and biochemical fractionation. Protein interactions were investigated using co-immunoprecipitation and proteomic analyses. For *in vivo* validation, a murine model of colorectal liver metastasis was generated by intrasplenic injection of tumor cells, followed by systemic administration of chondroitin sulfate–targeted nanoparticles delivering PCSK9 siRNA every 5 days for 18 days.

**Results:** PCSK9 was consistently expressed in liver sinusoidal endothelial cells and displayed a predominant nuclear localization, which increased upon tumor-induced activation. Proteomic integration identified multiple candidate interacting proteins involved in metabolic and tumor-related pathways. Targeted nanoparticle-mediated delivery achieved efficient PCSK9 silencing in vitro. In vivo, endothelial-specific PCSK9 inhibition significantly reduced liver metastatic tumor burden compared with control groups, whereas free siRNA showed no significant effect.

**Conclusions:** PCSK9 exhibits a novel nuclear localization in liver sinusoidal endothelial cells and potentially interacts with proteins implicated in tumor mediated pathways. Selective inhibition of endothelial PCSK9 using targeted nanoparticles significantly reduces colorectal liver metastasis, highlighting a novel therapeutic strategy focused on the hepatic microenvironment.

**Impact and Implications:** This study provides mechanistic insight into how PCSK9 contributes to colorectal liver metastasis by identifying its novel nuclear localization and potential function in liver sinusoidal endothelial cells. These findings are important for researchers and clinicians seeking to understand microenvironment-driven metastasis and resistance to current therapies. The demonstration that endothelial-specific targeting of PCSK9 reduces metastatic burden suggests a new avenue for therapeutic development beyond systemic inhibition. Such strategies could be translated into precision nanomedicine approaches to improve outcomes in patients with metastatic colorectal cancer while minimizing off-target effects.

## Introduction

Cancer is one of the leading causes of mortality worldwide, accounting for almost 20 million new cases each year. Among all malignancies, colorectal cancer (CRC) ranks third in incidence and shows a steadily increasing trend, driven in part by dietary and lifestyle factors (Bray *et al*, 2024; Ye *et al*, 2020). The primary cause of death in CRC patients is metastasis, with the liver being the most frequent metastatic site due to its unique anatomical and physiological characteristics. (Riihimäki *et al*, 2016; Vidal-Vanaclocha F., 2011)

Hepatic metastasis is a multistep process that involves tumor cell dissemination from the primary tumor, intravasation, circulation, extravasation, migration within the hepatic parenchyma, and the subsequent formation of micro- and macrometastases (Clark *et al*, 2016). Throughout this process, tumor cells actively remodel the hepatic microenvironment, transforming it into a pro-tumorigenic niche. The liver contains several specialized cell types that contribute to metastatic progression, including hepatocytes, Kupffer cells, hepatic stellate cells, and liver sinusoidal endothelial cells (LSECs) (Vidal-Vanaclocha, 2008). Each of these cell populations plays a distinct role in liver metastasis. For example, hepatic stellate cells promote liver metastasis through ANGPTL4 expression, which increases liver fibrosis, while Kupffer cells, the liver’s resident macrophages, can facilitate metastatic seeding and outgrowth through the induction of neutrophil extracellular trap (NET) formation (Fan *et al*, 2025; Tian *et al*, 2025).

Of particular interest in the metastatic cascade is the role of liver sinusoidal endothelial cells. LSECs are critically involved in tumor cell adhesion through the expression of adhesion molecules such as ICAM-1 and contribute to immune evasion via the expression of immune checkpoint molecules, including PD-1 (Benedicto *et al*, 2019; Shigeta *et al*, 2020; Carambia *et al*, 2014). Moreover, LSECs promote epithelial–mesenchymal transition (EMT) and angiogenesis, a process that is essential for the transition from micrometastases to macrometastases (Ou *et al*, 2014).

In this context, protein proprotein convertase subtilisin/kexin type 9 (PCSK9) has emerged as a molecule of considerable interest. PCSK9 is a member of the proprotein convertase family and is best known for its role in cholesterol homeostasis through the degradation of the low-density lipoprotein receptor (LDLR) (Lagace, 2014). This function has led to the development of monoclonal antibodies and siRNA-based therapies targeting PCSK9 for the treatment of atherosclerosis (Saad Cleto *et al*, 2026; Ray *et al*, 2023). However, PCSK9 has recently gained attention in oncology. Several studies have linked PCSK9 to tumor immunogenicity, as it has been shown to promote the degradation of major histocompatibility complex class I (MHC-I) molecules on tumor cells, thereby impairing antitumor immune responses (Liu *et al*, 2020; Singh *et al*, 2025). In addition, PCSK9 has been associated with the promotion of EMT by increasing the expression of Snail-1, N-cadherin, and MMP-9, while reducing E-cadherin levels (Wang *et al*, 2022). PCSK9 has also been linked to KRAS-driven signaling in colorectal carcinoma, contributing to a more aggressive tumor phenotype (Wong *et al*, 2022).

Beyond its role in tumor cells, PCSK9 has been implicated in shaping the tumor microenvironment. PCSK9 influences macrophage polarization, favoring a pro-tumorigenic phenotype, and has been shown to play a specific role in the hepatic vasculature (Hu *et al*, 2022). Notably, Gantz et al. demonstrated that PCSK9 promotes resistance to anti-angiogenic therapies in colorectal cancer liver metastases by facilitating vascular co-option, thereby reducing angiogenesis and leading to therapeutic failure (Gantz *et al*, 2025). Furthermore, colorectal cancer stem cells have been shown to induce a significant upregulation of PCSK9 expression in LSECs, enhancing their proliferative and migratory capacities (Martin *et al*, 2025). These findings highlight PCSK9 as a key regulator of LSEC function during liver metastasis.

Although several therapeutic strategies have been developed to inhibit PCSK9, current approaches result either in systemic inhibition, as in the case of monoclonal antibodies, or in hepatocyte-restricted targeting, such as siRNA-based therapies. Therefore, there is a compelling need to develop cell type–specific therapeutic strategies that enable selective inhibition of PCSK9 in key components of the metastatic niche. Particularly, targeting PCSK9 in LSECs may represent a promising approach given their central role in metastatic progression and the strong influence of PCSK9 on their pro-metastatic functions. Such strategies could lead to novel therapies aimed at reducing liver metastasis and improving the efficacy of anti-angiogenic treatments by limiting vascular co-option.

The aim of this study is to verify the expression of PCSK9 in LSECs and to develop a method for targeting PCSK9 in these cells using a cell-specific nanoparticle.

## Results

### 1. PCSK9 is present in LSEC after tumor-mediated activation

To investigate the effect of PCSK9 on liver sinusoidal endothelial cells (LSECs) during colorectal liver metastasis, conditioned media were obtained from differentiated human and murine metastatic colorectal tumor cells (SW620 and CT26) and colorectal cancer stem cells (CSCs)-like populations derived from SW620 (SW620-CSC) and CT26 (CT26-CSC) cells and used to stimulate human LSEC (hLSEC) cultures, as shown in Figure 1A. Immunofluorescence staining for PCSK9 and tubulin was subsequently performed on activated hLSEC cultures to assess both PCSK9 expression and its subcellular localization. PCSK9 was detected under all three experimental conditions and exhibited an apparently nuclear localization, as shown in Figure 1B. Similarly, Supplementary Figure 1A shows that the expression pattern observed in human endothelial cells is also present in murine endothelial cells. Notably, Supplementary Figure 1B reveals that PCSK9 displays a nuclear localization exclusively in hLSECs, which are responsible for the hepatic microvasculature, whereas in human umbilical vein endothelial cells HUVECs—derived from large blood vessels—PCSK9 is localized in the cytoplasm. This observation suggests that the nuclear localization of PCSK9 may be specific to endothelial cells of the microvasculature. To confirm this subcellular localization and to verify PCSK9 expression in human LSECs in vivo, immunostaining for CD31, an endothelial cell marker, together with PCSK9 was performed on liver tissue samples from colorectal cancer patients with liver metastases. These analyses demonstrated that PCSK9 is expressed in patient-derived LSECs and similarly exhibits a nuclear localization, as shown in Figure 1C.

**Figure 1.**
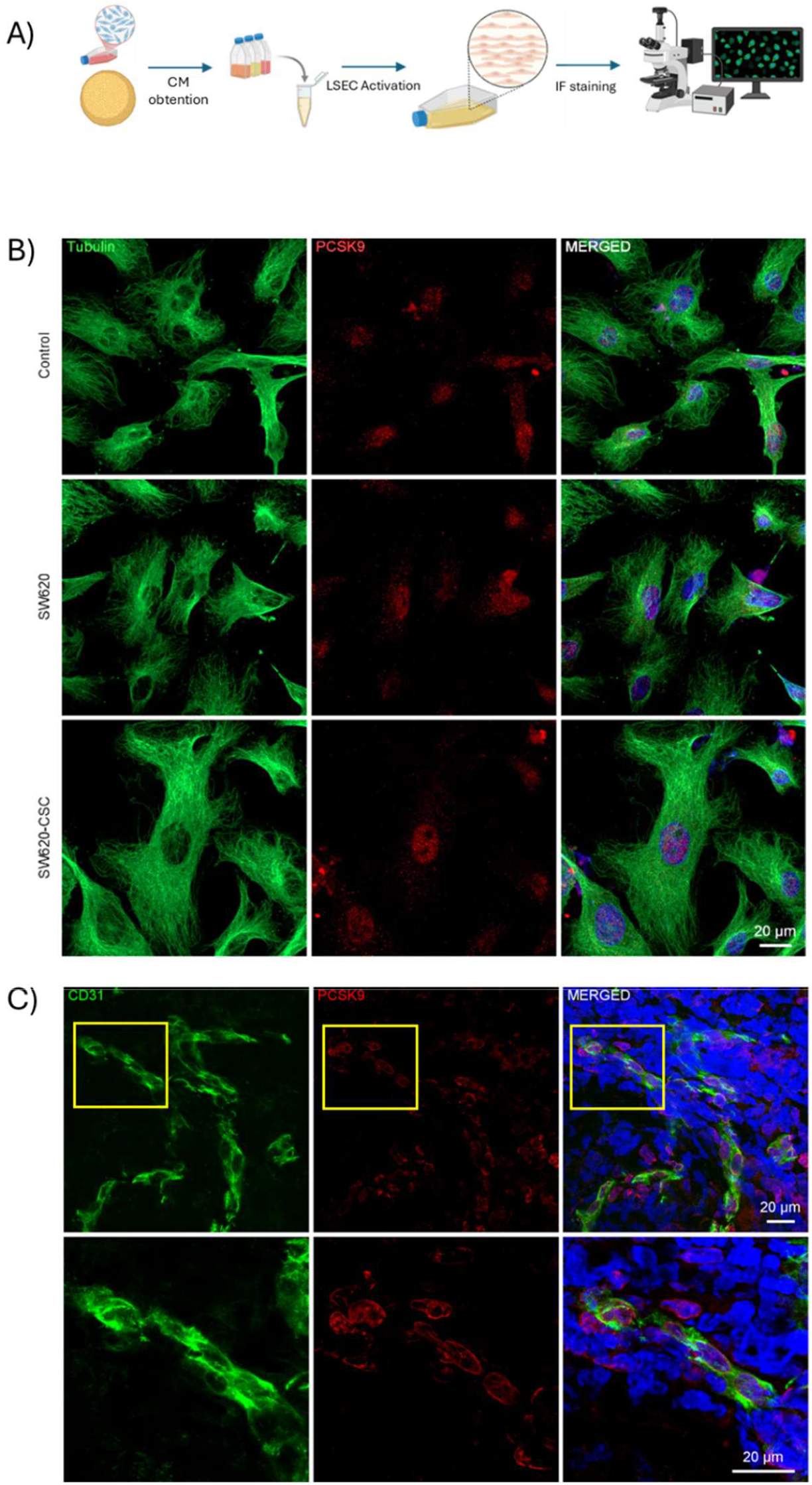
PCSK9 expression in human LSEC assessment. A) LSEC cell culture activation with differentiated metastatic colorectal tumor cells and colorectal CSC conditioned media. B) Immunofluorescent staining of PCSK9 (red), tubulin (green) and DAPI (blue) in human LSEC cell cultures in basal conditions and activation with differentiated metastatic colorectal tumor cells SW620 and colorectal CSC-like from SW620 conditioned media. C) DAPI (blue), CD31(green) and PCSK9 (red) staining on liver tissue samples from colorectal cancer patients with liver metastases. The areas highlighted in the images in the upper row are shown enlarged in the lower row.

### 2. PCSK9 is located in the nucleus of human LSEC

To investigate the subcellular localization of PCSK9, immunofluorescence staining was performed in cultures of hLSEC cell line using markers corresponding to the subcellular compartments in which PCSK9 has been previously reported, as well as a nuclear marker. Specifically, calnexin was used as an endoplasmic reticulum marker, TOM20 as a mitochondrial marker, LAMP1 as a lysosomal marker, and FUS/TLS as a nucleoplasmic marker, as shown in Figure 2A. Among these markers, the highest degree of colocalization was observed with FUS/TLS, suggesting that PCSK9 is localized in the nucleoplasm of these endothelial cells.

**Figure 2.**
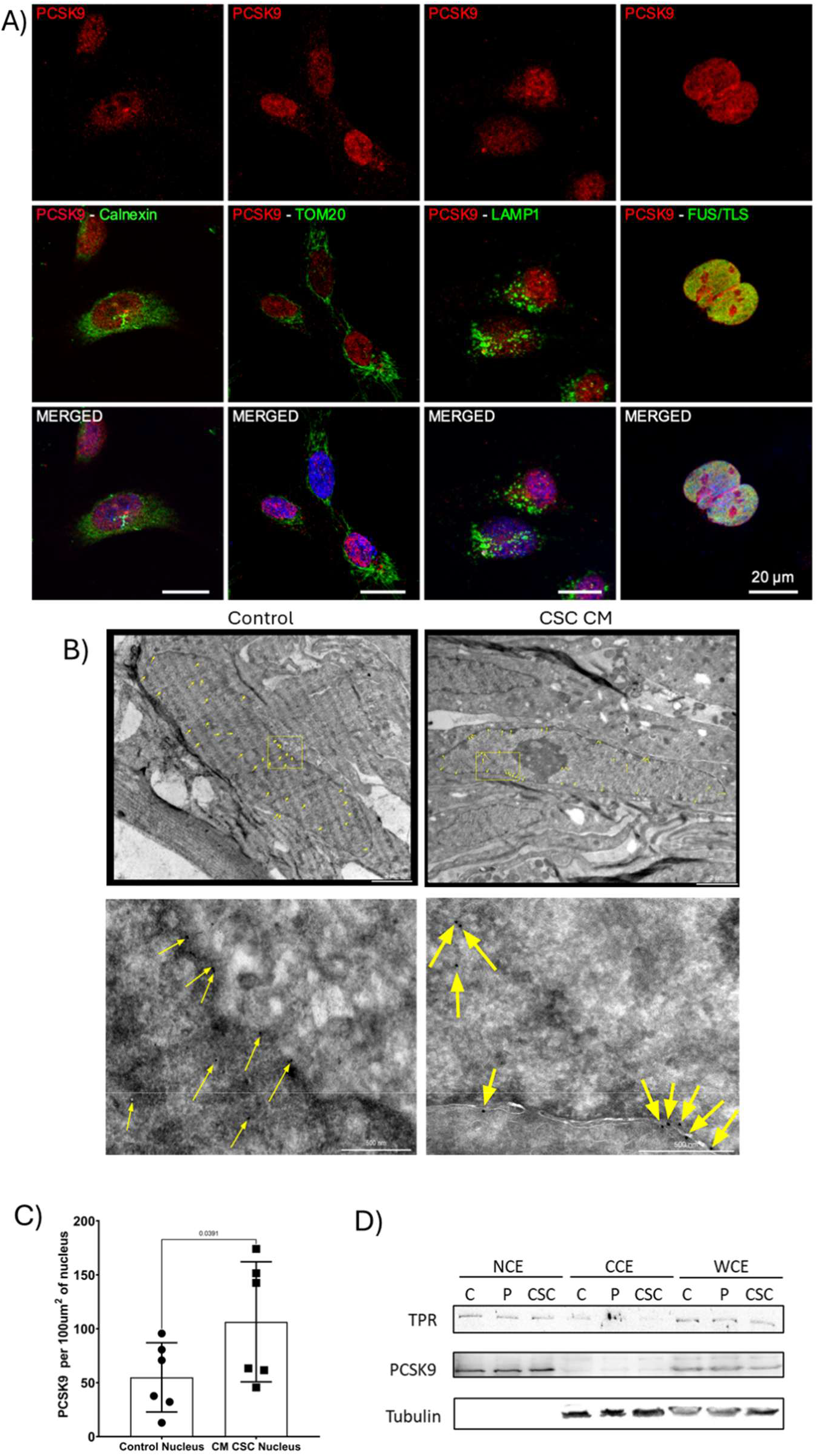
PCSK9 displays an intracellular and nuclear localization in human LSEC cultures. A) Immunofluorescence staining of PCSK9, DAPI and subcellular markers: calnexin (endoplasmic reticulum), TOM20 (mitochondria), LAMP1 (lysosomes), and FUS/TLS (nucleoplasm). B) Immunogold labeling of PCSK9 in control endothelial cells and cells activated with CSC-conditioned medium. The areas highlighted in the images in the upper row are shown enlarged in the lower row. C) Quantification of immunogold labeling in control and CSC-conditioned medium–activated endothelial cells Unpaired t-test. D) Western blot analysis of whole-cell extracts (WCE), cytoplasmic fractions (CCE), and nuclear fractions (NCE) from control and activated endothelial cells treated with conditioned media from differentiated tumor cells and cancer stem cells. Tubulin was used as a cytoplasmic marker and TPR as a nuclear marker.

To further confirm the intracellular localization of PCSK9, immunogold labeling was performed in basal hLSEC cultures and in hLSECs activated with SW620-CSC-conditioned media. As shown in Figure 2B, under both conditions the majority of PCSK9 signal was detected on the inner side of the nuclear envelope, confirming its intracellular and nuclear localization in this cell type. Moreover, quantification of the immunogold labeling (Figure 2C) revealed a higher density of PCSK9 per nuclear area in hLSECs activated with SW620-CSC-conditioned medium, suggesting that PCSK9 may play a role in tumor-induced activation of LSECs. Finally, Western blot analysis of nuclear extracts from control and activated hLSECs (Figure 2D) demonstrated that PCSK9 is predominantly enriched in the nuclear fraction, further confirming its intracellular localization in hLSECs.

### 3. PCSK9 Co-immunoprecipitation in HEK293 cells and LSEC proteomic study reveals PCSK9 potential interactors in LSEC

To investigate PCSK9 protein–protein interactions, a co-immunoprecipitation (co-IP) approach was performed. HEK293 cells were stably transfected with wild type PCSK9 (Supplementary Fig. 2A), and conditioned media were collected. PCSK9 was subsequently immunoprecipitated to identify its interacting partners by mass spectrometry analysis. The presence of PCSK9 in all samples was confirmed by Western blot analysis (Supplementary Fig. 2B).

This approach identified a large number of potential PCSK9-interacting proteins (Fig. 3A). Since the co-IP was performed in HEK293 cells, total proteomic analysis was conducted in control and activated LSEC cells to assess the relevance of these interactions in a more physiologically relevant context. Integration of both datasets revealed eight proteins common to the co-IP and LSEC proteomics (Fig. 3B). These proteins—pyruvate kinase, β-tubulin, spermine synthase, macrophage migration inhibitory factor, S100A11, dermcidin, fibulin-1, and follistatin-related protein 1—are shown in Fig. 3C and may represent candidate PCSK9 interactors in LSEC cells.

**Figure 3.**
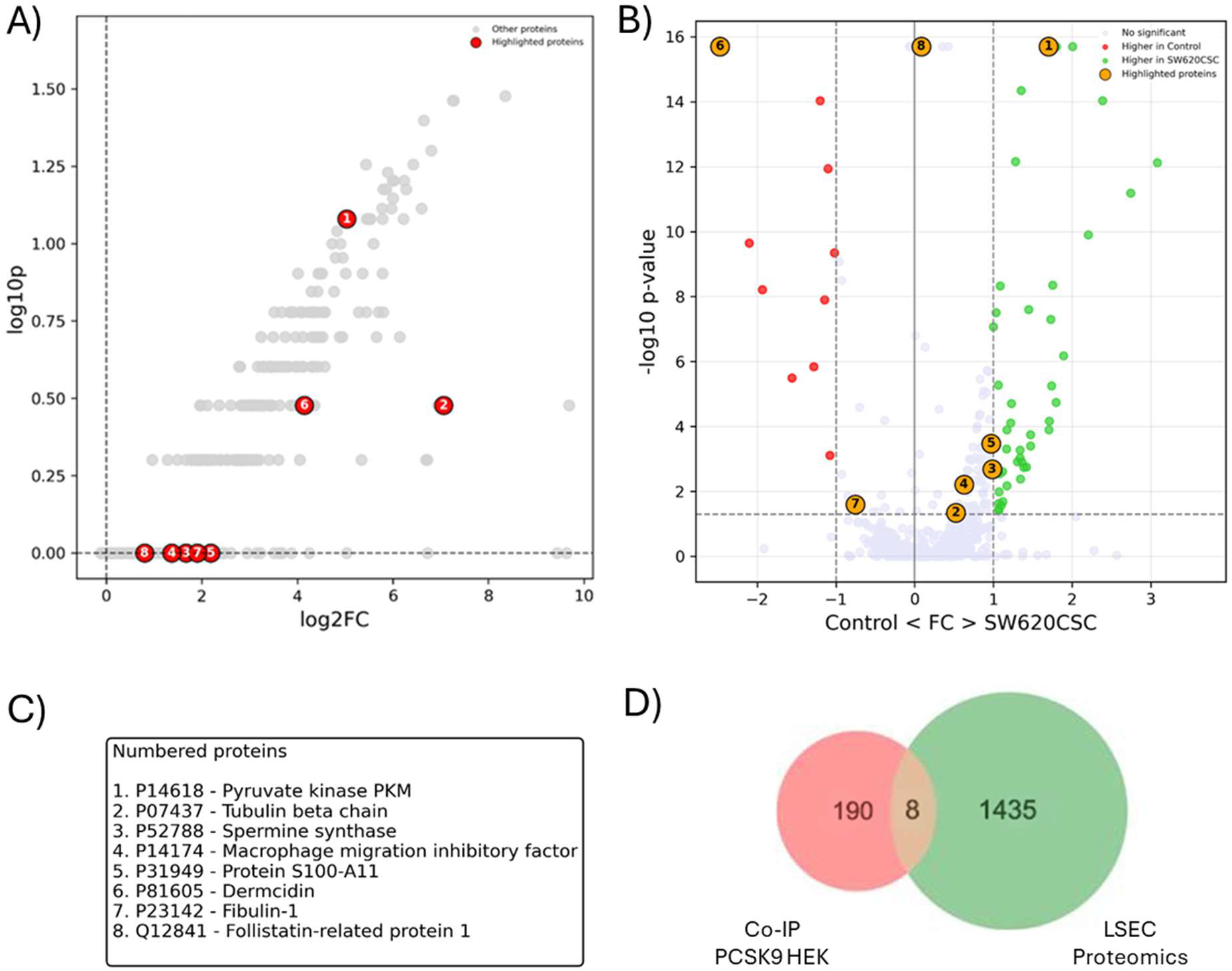
Identification of PCSK9 interactors in HEK293 cells and comparative analysis with proteomics of control and activated LSECs. A) Protein profile obtained from PCSK9 co-immunoprecipitation performed on conditioned media from HEK293 cells overexpressing PCSK9. Eight proteins of interest are highlighted. B) Comparative analysis of total proteomics from control and activated LSECs treated with SW620CSC conditioned media. The eight proteins identified in both the LSEC proteomics and the co-IP dataset are highlighted. C) Table showing proteins common to both datasets. D) Overview of the total number of proteins identified in each dataset and the overlap between them.

Overall, 198 proteins were identified in the co-IP and 1,443 in the total proteomic analysis, with eight proteins overlapping between both datasets (Fig. 3D).

### 4. PCSK9 can be inhibited in vitro in human LSEC using a chondroitin sulphate-conjugated sorbitan ester based nanoparticles loaded with selected siRNA

To specifically inhibit PCSK9 in endothelial cells, a nanoparticle previously developed by our group for the targeted delivery of miRNAs to hepatic endothelial cells was adapted for the encapsulation of PCSK9 siRNA (Marquez *et al*, 2018). This nanoparticle is composed of oleylamine and Span 80, which form the structural core of the nanoparticle, and chondroitin sulfate, which confers specificity toward LSECs. In addition, PCSK9 siRNA was incorporated into the nanoparticle; due to its electrical charge, the siRNA is retained within the inner compartment of the nanoparticle. The schematic structure of the nanoparticle is shown in Figure 4A.

**Figure 4.**
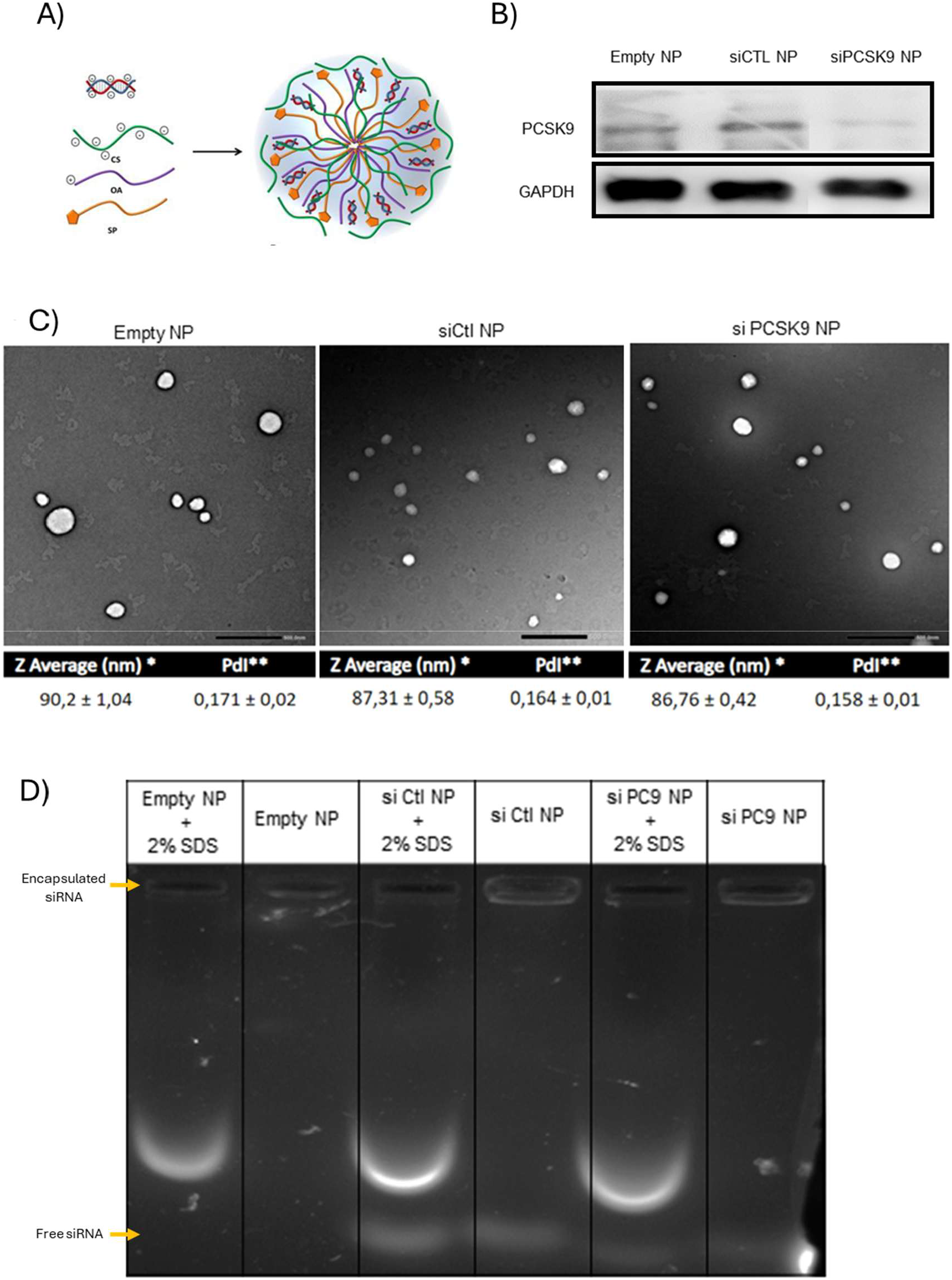
Nanoparticle characterization. A) Schematic representation of the nanoparticle used, showing the encapsulated siRNA, oleylamine (purple), Span 80 (orange), and chondroitin sulfate (green). B) Western blot analysis of PCSK9 expression in the Empty NP, siCtl NP, and siPCSK9 NP conditions 5 days after transfection; GAPDH was used as a housekeeping control. C) Representative uranyl acetate negative staining transmission electron microscopy (TEM) images of the Empty NP, siCtl NP, and siPCSK9 NP, together with dynamic light scattering (DLS) measurements. The Z-average value is shown as an indicator of mean nanoparticle size, and the polydispersity index (PDI) as a measure of nanoparticle homogeneity. D) Agarose gel (2%) electrophoresis of the three nanoparticle formulations incubated with or without 2% SDS. SDS treatment disrupts the nanoparticle structure, releasing the encapsulated nucleic acids and allowing their migration, thereby enabling assessment of nucleic acid localization within the nanoparticles. Arrows show encapsulated siRNA and Free siRNA.

To evaluate whether this nanoparticle effectively mediates PCSK9 silencing, murine LSEC cultures were incubated with the nanoparticle, as its use has been previously validated in murine models. Cells were treated with empty nanoparticles (Empty NP), nanoparticles containing a control siRNA (siCtl NP), or nanoparticles encapsulating PCSK9 siRNA (siPCSK9 NP). As shown by the Western blot analysis in Figure 4B, PCSK9 protein levels were markedly reduced five days after incubation with the siPCSK9 NP, whereas PCSK9 expression remained unchanged in the other two conditions.

After confirming the silencing efficiency of the functionalized nanoparticle, its size and siRNA encapsulation capacity were characterized. Transmission electron microscopy (TEM) images of Empty NP, siCtl NP, and siPCSK9 NP are shown in Figure 4C, demonstrating that nanoparticle integrity is preserved in all conditions. In addition, dynamic light scattering measurements revealed that all three nanoparticle formulations display diameters below 100 nm, indicating the generation of small-sized nanoparticles suitable for biological applications.

With respect to encapsulation efficiency, agarose gel electrophoresis analysis (Figure 4D) showed that nanoparticles treated with SDS (disrupted nanoparticles) migrated into the gel, whereas intact nanoparticles did not. Accordingly, intact siCtl NP and siPCSK9 NP emitted fluorescence only at the loading wells, indicating that the encapsulated nucleic acids were unable to migrate through the gel. This result confirms that a substantial fraction of the siRNA is successfully encapsulated within the nanoparticles, thereby validating this nanoparticle-based method for siRNA delivery.

### 5. Specific inhibition of LSEC PCSK9 in vivo significantly reduces liver colorectal metastatic tumor size

Once the quality of the nanoparticle was validated, *in vivo* experiments were performed. Colorectal tumor cells were injected into the spleen of mice, and livers were harvested for analysis 18 days later. As shown in Figure 5A, nanoparticles were administered every five days via tail vein injection, starting on day 1. The dosing schedule was selected based on the *in vitro* data shown in Figure 4B, which demonstrated that PCSK9 inhibition persisted for up to five days following nanoparticle treatment. To ensure the quality and consistency of the injected nanoparticles, particle size was measured by dynamic light scattering (DLS) for each nanoparticle preparation administered on each injection day, as detailed in Supplementary Table 1.

**Figure 5.**
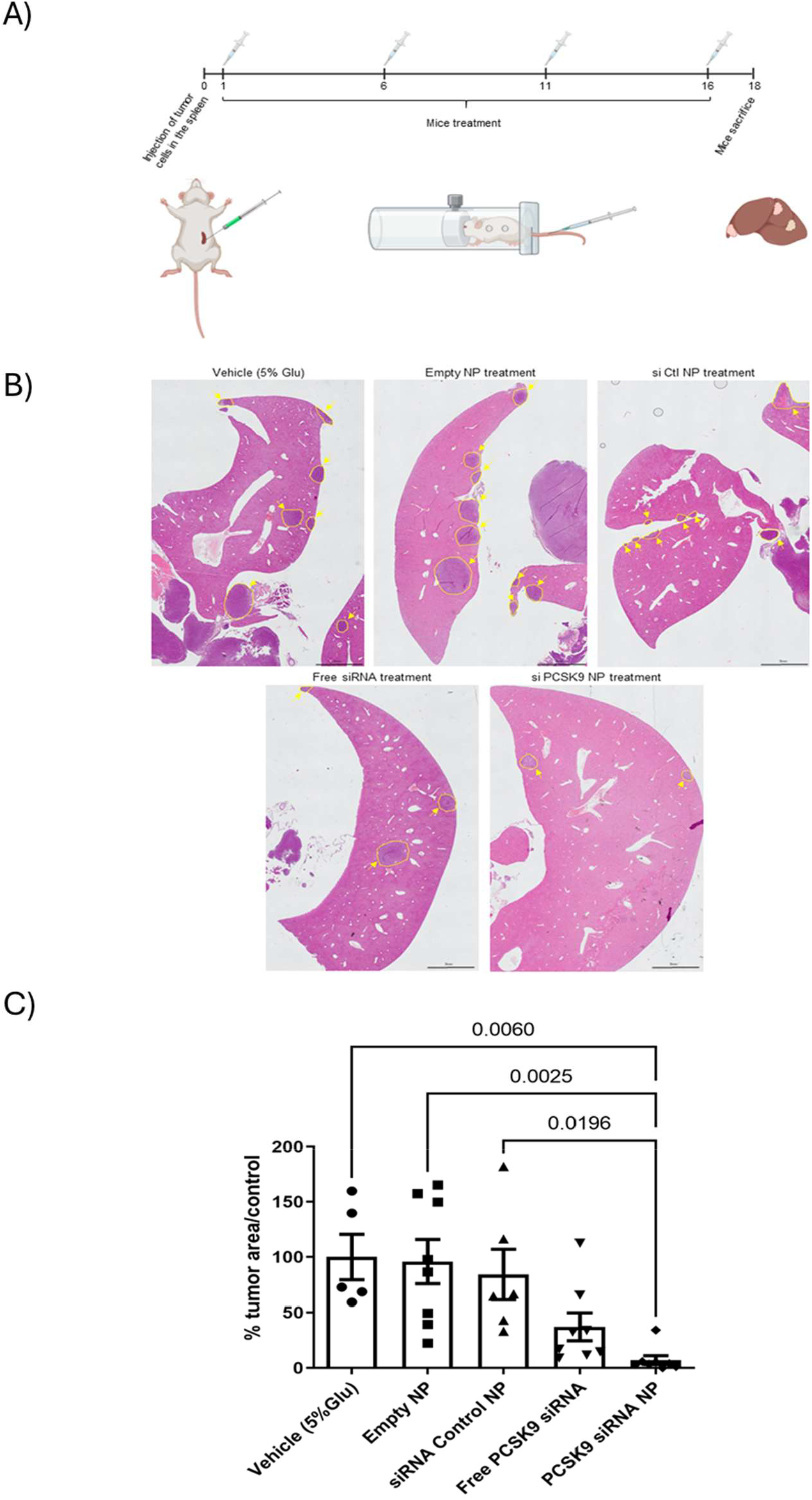
Endothelial PCSK9 inhibition *in vivo* significantly reduces colorectal liver metastatic tumor burden. A) Schematic representation of the experimental model. Colorectal tumor cells were injected into the spleen, nanoparticles were administered every five days via tail vein injection starting on day 1, and mice were sacrificed on day 18 for the analysis of liver metastases. B) Representative hematoxylin and eosin–stained liver sections from the in vivo experiments. Five experimental conditions are shown: vehicle (5% glucose), Empty NP, siCtl NP, free siRNA, and siPCSK9 NP. C) Quantification of tumor burden expressed as the percentage of tumor area normalized to the control condition. Comparisons among the five experimental groups are shown. Statistical analysis was performed using one-way ANOVA followed by Tukey’s post hoc test: ns, p > 0.05; *p < 0.05; **p < 0.01.

The *in vivo* results are presented in Figures 5B and 5C. Figure 5B shows representative hematoxylin and eosin–stained liver sections from the different experimental groups, with tumor-infiltrated areas highlighted in yellow in each field. The quantification of tumor burden is shown in Figure 5C. Upon normalization of tumor area percentages to the control condition, treatment with empty nanoparticles did not show any effect on tumor development. Similarly, treatment with control siRNA–loaded nanoparticles resulted in a slight, but non-significant, reduction in tumor area. Administration of free siRNA led to a marked decrease in tumor burden that, however, did not reach statistical significance. In contrast, mice treated with nanoparticles loaded with PCSK9 siRNA exhibited a significant reduction in tumor size compared with the control condition, the empty nanoparticle condition, and the control siRNA nanoparticle condition.

Altogether, these results demonstrate that specific inhibition of PCSK9 in LSECs using an endothelial-targeted nanoparticle loaded with PCSK9 siRNA significantly reduces colorectal liver metastasis in an in vivo model.

## Discussion

PCSK9 was originally identified as a central regulator of cholesterol homeostasis through its role in LDL receptor degradation (Benjannet *et al*, 2004). However, beyond its well-established metabolic function, increasing evidence supports a role for PCSK9 in cancer progression. Elevated PCSK9 expression has been associated with poor prognosis in multiple tumor types, including colorectal cancer, particularly in the metastatic setting (Bassi *et al*, 2017; Gantz *et al*, 2025; Scamuffa *et al*, 2008; Sun *et al*, 2012). Notably, most studies addressing the role of PCSK9 in cancer have focused on tumor cells themselves, with limited attention paid to its function within the tumor microenvironment. Given the key contribution of hepatic stromal cells to colorectal liver metastasis, understanding the role of PCSK9 in non-tumoral compartments, such as liver sinusoidal endothelial cells (LSECs), is of relevance.

In this study, we investigated the role of PCSK9 in LSECs during colorectal liver metastasis, focusing on its subcellular localization and functional relevance *in vivo*. To model tumor-driven endothelial activation, we used colorectal cancer stem cell (CSC)–conditioned media. This approach was chosen because CSCs have been reported to overexpress PCSK9 and because endothelial activation and mobilization are initiated early during metastatic seeding, coinciding with the appearance of CSCs in the metastatic niche (Martin *et al*, 2025). Accordingly, CSC-conditioned media provides a biologically relevant stimulus that mirrors early microenvironmental changes occurring during metastatic colonization.

We demonstrate that PCSK9 is expressed by both human and murine LSEC and is present in this cell type within liver tissue from patients with colorectal liver metastases. A key and novel finding of our study is the predominant nuclear localization of PCSK9 in LSECs. This observation was consistently supported by multiple complementary approaches, including immunofluorescence with subcellular markers, immunogold electron microscopy, and biochemical fractionation of cytoplasmic and nuclear compartments. Importantly, nuclear PCSK9 levels were increased in LSECs activated with CSC-conditioned media, suggesting that tumor-derived signals not only induce PCSK9 expression but also influence its intracellular distribution.

Previous work by Martín et al. described increased PCSK9 mRNA expression in activated LSECs; however, the nuclear localization identified here represents a novel aspect of PCSK9 biology (Martin *et al*, 2025). While PCSK9 is classically considered a secreted protein acting through receptor trafficking, it has also been shown to function as a carrier for intracellular proteins such as LDLR and MHC-I, directing them toward lysosomal degradation (Liu *et al*, 2020; Lagace, 2014). These observations raise the possibility that nuclear PCSK9 may interact with specific nuclear or nuclear-associated proteins in LSECs, thereby contributing to endothelial activation and tumor progression.

To explore this hypothesis, we combined PCSK9 co-immunoprecipitation with proteomic profiling of control and CSC-activated LSECs. By intersecting these datasets, we identified a subset of eight proteins that interact with PCSK9 and are detected in the proteomic analysis of basal and activated LSECs. Among these, five proteins—pyruvate kinase PKM, β-tubulin, macrophage migration inhibitory factor (MIF), spermine synthase, and S100A11—were upregulated following LSEC activation with CSC-conditioned media. Notably, PKM was the only protein significantly overexpressed in LSEC activated with CSC-conditioned media, whereas the other proteins exhibited higher expression levels that did not reach statistical significance. In addition, PKM, MIF, S100A11, and spermine synthase have been reported to localize the nucleus and to play roles in cancer progression.

The PKM2 isoform is of particular interest, as it has been described as a regulator of colorectal liver metastasis in colorectal cancer cells. For example, KHK-A–mediated phosphorylation of PKM2 promotes its nuclear translocation, thereby enhancing the metastatic potential of tumor cells (Peng *et al*, 2024). In addition, in hypoxic pancreatic tumors, nuclear PKM2 interferes with NF-κB/p65 and HIF-1α signaling, leading to VEGF-A secretion and subsequent angiogenesis, highlighting PKM2 as a relevant target to study in the context of PCSK9 interaction (Azoitei *et al*, 2016).

MIF has been shown to promote colorectal liver metastasis when secreted by LSECs; however, it has also been reported to translocate to the nucleus, where it can exert nuclease activity (Hu *et al*, 2015; Xiao *et al*, 2022). Therefore, the interaction between PCSK9 and MIF may be related to this nuclear function of MIF in endothelial cells. Similarly, S100A11 has been reported to promote metastasis via activation of AKT and ERK signaling pathways in hepatocellular carcinoma. This effect is mediated by changes in the nuclear-to-cytoplasmic ratio of S100A11 in response to intracellular calcium levels, a mechanism that could explain a nuclear interaction with PCSK9 (Zheng *et al*, 2023). Finally, spermine synthase has been implicated in liver macrophage polarization and in the suppression of antitumor immunity in hepatocellular carcinoma through the PI3K–AKT–mTOR–S6K pathway (Sun *et al*, 2025). Importantly, spermine synthase also participates in epigenetic regulation by controlling nuclear polyamine levels required for histone acetylation (Iacomino *et al*, 2012).

Taken together, these interactions provide a plausible mechanistic explanation for the nuclear accumulation of PCSK9 and support a role for PCSK9 as a modulator of endothelial pathways linked to tumor growth and metastatic progression.

Although *in vitro* studies have shown that PCSK9 inhibition reduces LSEC proliferation and migration, its endothelial-specific role *in vivo* has remained insufficiently defined (Martin *et al*, 2025). Previous work demonstrated that systemic inhibition of PCSK9 enhances anti-angiogenic therapies by limiting vascular co-option (Gantz *et al*, 2025). In contrast, our study specifically targeted PCSK9 in LSECs using a lipidic nanoparticle optimized for endothelial delivery (Marquez *et al*, 2018). We show that this targeted approach results in a significant reduction in colorectal liver metastasis burden, whereas systemic administration of free PCSK9 siRNA produced only a non-significant effect. These findings underscore the importance of cell-specific targeting and highlight LSECs as a critical compartment through which PCSK9 modulates metastatic progression.

Mechanistically, the observed reduction in metastatic burden could be related to angiogenic and endothelial activation processes. Angiogenesis is required for the transition from micrometastases to macrometastases, and LSECs play a central role in this process (Cañellas-Socias *et al*, 2022). By limiting PCSK9 activity in LSECs, tumor-induced endothelial remodeling may be attenuated, thereby constraining metastatic outgrowth.

In conclusion, our study identifies PCSK9 as a nuclear protein in LSECs, uncovers its association with tumor-promoting endothelial pathways, and demonstrates that endothelial-specific PCSK9 inhibition significantly reduces colorectal liver metastasis *in vivo*. These findings expand the functional landscape of PCSK9 beyond lipid metabolism and position it as a novel regulator of the hepatic endothelial microenvironment. Future studies will be required to elucidate the mechanisms underlying PCSK9 nuclear translocation and to functionally validate its interactions with nuclear proteins in LSECs, with the goal of exploiting PCSK9 as a therapeutic target in metastatic liver disease.

## Materials and Methods

### Cell Culture and Cancer Stem Cell Derivation

Immortalized liver sinusoidal endothelial cells (LSECs) from both, human (hLSEC) and murine (mLSEC) origins were obtained from Innoprot (Innoprot). These cells were cultured as adherent monolayers on culture plates pretreated with 10% rat-tail type I collagen (08-115, Sigma Aldrich) for 1 hour at room temperature, at 37ºC and 5% CO2. Complete Endothelial Cell Medium (P60104, Innoprot) was used to maintain the cultures. HUVEC cells (C0035C, Thermo Fisher) were cultured as adherent monolayers in treated culture plates precoated with Gelatine (G1393, Merck) for 2 hours at room temperature. Endothelial Cell Growth Medium (C-22110, Sigma-Aldrich) supplemented with 2% Fetal Bovine Serum (C-22110, Sigma-Aldrich), 1% Endothelial Cell Growth Supplement (C-22110, Sigma-Aldrich) and 100µg/ml Penicillin/Streptomycin solution (C-22110, Sigma-Aldrich) were used for cell culture. The human metastatic colorectal adenocarcinoma cell line SW620 and the murine metastatic colorectal adenocarcinoma cell line CT26 were acquired from the American Type Culture Collection (ATCC). SW620 cells were maintained in Leibovitz’s L15 medium (L5520, Sigma Aldrich), while CT26 cells were cultured in fully supplemented RPMI medium (R8758, Sigma Aldrich). To derive cancer stem cell (CSC)-like populations from SW620 and CT26 cells, 1 × 10^4^ cells per well were seeded in low-adherence 6-well plates and cultured for 7 days in serum-free DMEM-F12 medium (11554546, GIBCO). The medium was supplemented with 1% v/v N2, 0.5% v/v B27, 20 ng/mL fibroblast growth factor (FGF), and 20 ng/mL epidermal growth factor (EGF). Conditioned media were obtained after incubation of cell cultures during 24 hours with non-complete Endothelial Cell Media. Conditioned media was diluted (1:1 v/v) with complete media before being added to the cells. To obtain the condition activated with CSC conditioned media in absence of PCSK9, LSEC were pretreated 24 hours before stimulation with 50 µg/mL PCSK9 chemical inhibitor to inhibit PCSK9 translation (PF-06446846 hydrochloride, Sigma-Aldrich).

### Protein Extraction and Western Blot

LSECs were washed twice with cold 1X PBS and subsequently lysed using RIPA Buffer at a concentration of 100 µL per 106 cells. Protein concentrations were determined using the Bicinchoninic Acid (BCA) assay (Sigma-Aldrich). Samples were resuspended in 4X loading buffer (250mM Tris-HCl pH 6.8, 500mM β-mercaptoethanol, 50% glycerol, 10% SDS and bromophenol blue). For each sample, 20 µg of protein lysate was loaded onto a 10% SDS-PAGE gel and electrophoresed for 1 hour and 30 minutes at 90 V. Proteins were then transferred onto a nitrocellulose membrane under wet transfer conditions at a constant current of 385 mA for 3 hours and 30 minutes, with cooling on ice. Following protein transfer, membranes were incubated overnight at 4°C with primary antibodies, including rabbit anti-PCSK9 (1:2000, ab181142, Abcam), mouse anti-αtubulin (1:1000, T5168, Sigma-Aldrich) and anti-TPR (1:2000, abcam ab84516). Subsequently, the membranes were incubated with horseradish peroxidase (HRP)-conjugated secondary antibodies (anti-mouse HRP (ab6728, Abcam) and anti-rabbit HRP (ab102279, Abcam) for 1 hour and 30 minutes at room temperature. Protein detection was performed using Immobilon Crescendo Western HRP substrate (WBLUR0100, Merck), and chemiluminescence signals were visualized and captured using the iBright CL1500 Imaging System (Invitrogen, Thermo Fisher Scientific).

### Nuclear fractions isolation

Nuclear and cytoplasmic fractions were isolated using a modified REAP protocol (Suzuki et al, 2010). Cells were scraped, resuspended in PBS containing 0.2% IGEPAL, and homogenized by pipetting. An aliquot was collected as whole-cell lysate, while the remaining suspension was centrifuged at 200 g for 3 min to separate cytoplasmic (supernatant) and nuclear (pellet) fractions. The nuclear pellet was washed once and lysed in 1X Laemmli buffer. Cytoplasmic and whole-cell fractions were supplemented with 4× Laemmli buffer. All fractions were sonicated briefly to ensure complete nuclear disruption prior to analysis.

### Cryo-Immunogold Electron Microscopy

Cells were fixed with 4% paraformaldehyde and 0.1% glutaraldehyde for 15 min, washed in PBS, scraped, and pelleted. Samples were embedded in 12% gelatin, infiltrated with 2.3 M sucrose at 4 °C for 24 h, frozen in liquid nitrogen, and sectioned at −90 °C (50 nm) using a Leica Cryo-UC7. Cryosections were collected on formvar-coated nickel grids. After quenching with glycine and blocking, grids were incubated with an anti-PCSK9 primary antibody (1:50), followed by a 10-nm gold-conjugated donkey anti-goat secondary antibody (1:60). Sections were post-fixed with 2% glutaraldehyde, contrasted with uranyl acetate in methylcellulose, and imaged using a Hitachi H7650 transmission electron microscope. Image analysis was performed with ImageJ.

### Immunofluorescent Staining

LSEC cells were seeded on collagen-coated coverslips, fixed with 4% paraformaldehyde (PFA), and permeabilized with PBS-Triton X-100 and BSA. After blocking, cells were incubated overnight with primary antibodies, followed by fluorophore-conjugated secondary antibodies and DAPI for nuclear staining. Coverslips were mounted with Fluoromount-G, and imaging was performed using a Zeiss LSM800 confocal microscope with ImageJ software for processing. Regarding tissue inmunofluorecent staining, liver tissue sections were prepared using a cryostat and fixed with 4% PFA. After permeabilization and blocking, tissues were incubated with primary antibodies overnight, followed by secondary antibodies and DAPI. To reduce autofluorescence, tissues were treated with Sudan Black B and ethanol. Finally, slides were mounted with Fluoromount-G and stored at 4°C for long-term use. Used antibodies are shown in Table 1. Image processing was performed using Fiji Image J software. The representations appearing in the results were obtained using the Z-stack option of this software

### Primary antibodies

| Antibody | Dilution | Supplier | Reference |
| --- | --- | --- | --- |
| Goat polyclonal anti-PCSK9 | 1:200 | abcam | ab28770 |
| Mouse polyclonal anti-αTubulin | 1:1000 | Sigma-Aldrich | T5168 |
| Mouse monoclonal anti-TOM20 | 1:500 | Santa Cruz | Sc-17764 |
| Rabbit polyclonal anti-Calnexin | 1:500 | Santa Cruz | Sc-46669 |
| Mouse monoclonal anti-FUS/TLS | 1:500 | abcam | ab124923 |
| Mouse monoclonal anti-CD31 | 1:200 | DAKO | M0823 |

### PCSK9 overexpression, purification, and mass spectrometry analysis

HEK293 cells stably expressing the PCSK9 plasmid (pCMV-PCSK9-FLAG) were generated by selection with Geneticin (G418; Sigma-Aldrich). Cells were expanded to near confluence, washed, and incubated in serum-free medium (Opti-MEM or PBS). Conditioned media were collected, clarified by centrifugation, and incubated with Ni-NTA resin for 2 h at 4 °C. After washing, bound proteins were eluted with 500 mM imidazole. Eluted fractions were analyzed by SDS-PAGE and Coomassie staining, and PCSK9-containing fractions were pooled, quantified, supplemented with 10% glycerol, and stored.

For protein identification, PCSK9 bands were excised and subjected to in-gel tryptic digestion following a modified Shevchenko protocol. Peptides were extracted, desalted using C18 microcolumns, and analyzed by LC-MS/MS using an EASY-nLC 1000 system coupled to a Q Exactive mass spectrometer (Thermo Scientific). Peptides were separated on Acclaim PepMap columns with a linear 2–40% acetonitrile gradient. MS data were acquired in data-dependent mode.

Raw files were processed using Proteome Discoverer 2.2 and searched against the UniProt-SwissProt human database. Carbamidomethylation of cysteine was set as a fixed modification and methionine oxidation as a variable modification. Identified proteins were exported for downstream analysis.

### LSEC proteomic assay

Protein extracts from 1 × 10^6^ human liver sinusoidal endothelial cells (LSECs) were resolved by short migration on 10% SDS-PAGE gels and visualized by colloidal Coomassie staining. Bands of interest were excised, reduced, alkylated, and digested overnight with trypsin. Peptides were extracted, acidified, and analyzed by LC-MS/MS on an Orbitrap Lumos mass spectrometer using a 146-min gradient. Data were processed with Proteome Discoverer 2.5 (Sequest HT) against the UniProt human database, allowing up to two missed cleavages. Carbamidomethylation of cysteine was set as a fixed modification, while methionine oxidation, N-terminal acetylation, methionine loss, and methionine loss/acetylation were included as variable modifications. Protein grouping was performed based on peptide sequence homology to define master proteins, and known contaminants were annotated.

### Proteomic statistical analysis

A comparative analysis was performed using two independent proteomic datasets: a relative abundance dataset from LSEC cells comparing CSC and CTL conditions, and a PCSK9 immunoprecipitation dataset obtained from HEK cells. The objective was to identify proteins that were significantly altered in LSEC cells and also detected in the PCSK9 immunoprecipitate, suggesting potential association with PCSK9.

Data were processed using custom Python scripts based on pandas, NumPy and matplotlib. Protein identifiers were standardized by removing spaces, converting entries to uppercase and eliminating isoform suffixes after hyphens. Numerical values were cleaned by standardizing decimal separators and converting abundance and p-value columns into numeric format. In the LSEC dataset, proteins with p-values ≤ 0.05 were considered statistically significant.

The LSEC and immunoprecipitation datasets were merged according to their standardized protein accession identifiers. Proteins present in both datasets were classified as common proteins, and those that also met the statistical significance threshold in the LSEC comparison were classified as common significant proteins. Abundance values were transformed as logarithm and represented in scatter plots comparing CSC/CTL abundance with immunoprecipitation abundance. Common significant proteins were highlighted, and selected proteins of interest were numbered and annotated in the corresponding legends.

### Span-Based Nanoparticle Formulation

Span-based nanoparticles were prepared by ethanol injection. Span 80 and oleylamine were dissolved in ethanol (6.6 and 0.33 mg/mL, respectively), while chondroitin sulfate (0.125 mg/mL) and siRNA (500 nM) were dissolved in the aqueous phase. Nanoparticles formed spontaneously upon dropwise addition of the ethanolic phase into the aqueous phase (1:2, v/v) under magnetic stirring. Ethanol was removed and nanoparticle suspension concentrated nine-fold by rotary evaporation. Particle size and zeta potential were measured by dynamic light scattering (Zetasizer), morphology was assessed by transmission electron microscopy after uranyl acetate negative staining, and siRNA association efficiency was evaluated by 2% agarose gel electrophoresis.

### Cell culture transfection with the span-based nanoparticle

PCSK9-targeting siRNA (sense: 5′-GGGUCAUGGUCACCGACUU-3′; antisense: 5′-AAGUCGGUGACCAUGACCC-3′; Sigma-Aldrich) and non-targeting control siRNA (4390844, Thermo Fisher Scientific) were encapsulated in a span–based lipidic nanoparticle. LSECs were transfected with siPCSK9 or control siRNA at a final concentration of 10 nM. Briefly, 1 × 10^5^ cells were seeded in collagen I–coated 6-well plates and incubated overnight. Cells were then treated for 4 h with nanoparticles diluted in Opti-MEM and incomplete endothelial cell medium, after which fresh complete medium was added. PCSK9 knockdown was assessed by Western blot at day 5 post-transfection.

### Mouse Model of Colorectal Cancer Liver Metastasis

Liver metastases were induced by intrasplenic injection of CT26 colon carcinoma cells into Balb/c mice. Mice were anesthetized with isoflurane, and 1.5 × 10^5^ cells in 100 µL PBS were injected into the spleen, followed by surgical closure. Animals were randomly assigned to five treatment groups (glucose, empty nanoparticles, control siRNA nanoparticles, free PCSK9 siRNA, or PCSK9 siRNA nanoparticles) and treated by tail-vein injection every 5 days starting one day post-inoculation, with doses adjusted to body weight. Mice were sacrificed 18 days after tumor cell injection.

### Tissue Processing and Histological Analysis

Livers and spleens were fixed in 4% paraformaldehyde, dehydrated, paraffin-embedded, and sectioned (5 µm). Liver sections were stained with hematoxylin and eosin, scanned, and metastatic burden was quantified using FIJI by calculating tumor area relative to total liver area.

## Author contributions

A.M., A.-M.K. and I.B. (Iker Badiola) were responsible for the study concept and design. A.M., M.D.G.-E., N.F., I.B. (Iratxe Bañares), P.G.G., A.B.V., J.M. and H.G.G. performed the investigation and formal analysis. A.M. and I.B. contributed to the methodology and writing of the original draft. J.E. performed bioinformatic analysis. A.-M.K., A.S.B., O.C., P.G.G., F.J.S., and I.B. (Iker Badiola) handled conceptualization, funding acquisition and supervision. All authors have read and agreed to the published version of the manuscript.

## Disclosure and competing interests statement Nothing to declare

## Acknowledgements

We would like to thank Ricardo Andrade and Alex Díez from the Analytical and High-Resolution Microscopy Service in Biomedicine of the SGIker services (EHU), and the technician Melina Petrel of the Bordeaux Imaging Center (BIC). Mass spectrometry analysis was performed in the Proteomics Core Facility (SGIKER) at the University of the Basque Country and the CGFB Proteomics Core Facility at Bordeaux of University.

## References and citations

Azoitei N, Becher A, Steinestel K, Rouhi A, Diepold K, Genze F, Simmet T C Seufferlein T (2016) PKM2 promotes tumor angiogenesis by regulating HIF-1α through NF-κB activation. Molecular Cancer 2016 15:1 15: 3-

Bassi DE, Fu J, De Cicco RL, Klein-Szanto AJPAJP, Seidah NG, Prat A, Michel Chrétien, Seidah NG, Artenstein AW, Opal SM, et al (2017) Selective inhibition of proprotein convertases represses the metastatic potential of human colorectal tumor cells. Mol Carcinog 5: 1122–1131

Benedicto A, Herrero A, Romayor I, Marquez J, Smedsrød B, Olaso E & Arteta B (2019) Liver sinusoidal endothelial cell ICAM-1 mediated tumor/endothelial crosstalk drives the development of liver metastasis by initiating inflammatory and angiogenic responses. Sci Rep 9: 1–12

Benjannet S, Rhainds D, Essalmani R, Mayne J, Wickham L, Jin W, Asselin MC, Hamelin J, Varret M, Allard D, et al (2004) NARC-1/PCSK9 and its natural mutants: Zymogen cleavage and effects on the low density lipoprotein (LDL) receptor and LDL cholesterol. Journal of Biological Chemistry 279: 48865–48875

Bray F, Laversanne M, Sung H, Ferlay J, Siegel RL, Soerjomataram I & Jemal A (2024) Global cancer statistics 2022: GLOBOCAN estimates of incidence and mortality worldwide for 36 cancers in 185 countries. CA Cancer J Clin 74: 229–263

Cañellas-Socias A, Cortina C, Hernando-Momblona X, Palomo-Ponce S, Mulholland EJ, Turon G, Mateo L, Conti S, Roman O, Sevillano M, et al (2022) Metastatic recurrence in colorectal cancer arises from residual EMP1+ cells. Nature 611: 603–613

Carambia A, Freund B, Schwinge D, Heine M, Laschtowitz A, Huber S, Wraith DC, Korn T, Schramm C, Lohse AW, et al (2014) TGF-β-dependent induction of CD4+CD25+Foxp3+ Tregs by liver sinusoidal endothelial cells. J Hepatol 61: 594–599

Clark AM, Ma B, Taylor DL, Griffith L C Wells A (2016) Liver metastases: Microenvironments and ex-vivo models. Exp Biol Med 241: 1639–1652

Fan X, Li B, Zhang F, Liu M, Kwan HY, Liu Z & Su T (2025) FGF19-Activated Hepatic Stellate Cells Release ANGPTL4 that Promotes Colorectal Cancer Liver Metastasis. Adv Sci (Weinh) 12

Gantz A, Kapelanski-Lamoureux A, Rada M, Krzywon L, Tsamchoe M, Tsatoumas A, Petrillo SK, Seidah NG, Lazaris A & Metrakos P (2025) High levels of serum cholesterol increase the risk of developing vessel co-opting tumors in colorectal cancer liver metastases. FEBS Journal

Hu CT, Guo LL, Feng N, Zhang L, Zhou N, Ma LL, Shen L, Tong GH, Yan QW, Zhu SJ, et al (2015) MIF, secreted by human hepatic sinusoidal endothelial cells, promotes chemotaxis and outgrowth of colorectal cancer in liver prometastasis. Oncotarget 6: 22410–22423

Hu J, Zhang M, Gui L, Wan Q, Zhong J, Bai L & He M (2022) PCSK9 Suppresses M2-Like Tumor-Associated Macrophage Polarization by Regulating the Secretion of OX40L from Hepatocellular Carcinoma Cells. Immunol Invest 51: 1678–1693

Iacomino G, Picariello G & D’Agostino L (2012) DNA and nuclear aggregates of polyamines. Biochimica et Biophysica Acta (BBA) - Molecular Cell Research 1823: 1745–1755

Lagace TA (2014) PCSK9 and LDLR degradation: Regulatory mechanisms in circulation and in cells. Curr Opin Lipidol 25: 387–393

Liu X, Bao X, Hu M, Chang H, Jiao M, Cheng J, Xie L, Huang Q, Li F & Li CY (2020) Inhibition of PCSK9 potentiates immune checkpoint therapy for cancer. Nature 2020 588:7839 588: 693–698

Marquez J, Fernandez-Piñeiro I, Araúzo-Bravo MJ, Poschmann G, Stühler K, Khatib AM, Sanchez A, Unda F, Ibarretxe G, Bernales I, et al (2018) Targeting liver sinusoidal endothelial cells with miR-20a-loaded nanoparticles reduces murine colon cancer metastasis to the liver. Int J Cancer 143: 709–719

Martin A, Gerovska D, Arauzo-Bravo MJ, Duarte García-Escudero M, García García H, Bañares I, Fontal N, Siegfried G, Evrad S, Pernot S, et al (2025) Inhibition of PCSK9 Attenuates Liver Endothelial Cell Activation Induced by Colorectal Cancer Stem Cells during Liver Metastasis. Cancers (Basel) 16

Ou J, Peng Y, Deng J, Miao H, Zhou J, Zha L, Zhou R, Yu L, Shi H & Liang H (2014) Endothelial cell-derived fibronectin extra domain A promotes colorectal cancer metastasis via inducing epithelial–mesenchymal transition. Carcinogenesis 35: 1661–1670

Peng C, Yang P, Zhang D, Jin C, Peng W, Wang T, Sun Q, Chen Z, Feng Y & Sun Y (2024) KHK-A promotes fructose-dependent colorectal cancer liver metastasis by facilitating the phosphorylation and translocation of PKM2. Acta Pharm Sin B 14: 2959–2976

Ray KK, Troquay RPT, Visseren FLJ, Leiter LA, Scott Wright R, Vikarunnessa S, Talloczy Z, Zang X, Maheux P, Lesogor A, et al (2023) Long-term efficacy and safety of inclisiran in patients with high cardiovascular risk and elevated LDL cholesterol (ORION-3): results from the 4-year open-label extension of the ORION-1 trial. Lancet Diabetes Endocrinol 11: 109–119

Riihimäki M, Hemminki A, Sundquist J & Hemminki K (2016) Patterns of metastasis in colon and rectal cancer OPEN. Nature Publishing Group

Saad Cleto A, Schirlo JM, Machozeki J & Martins CM (2026) Alirocumab versus Evolocumab on Cardiovascular Outcomes: A Systematic Review and Meta-analysis. Curr Cardiol Rev 22

Scamuffa N, Siegfried G, Bontemps Y, Ma L, Basak A, Cherel G, Calvo F, Seidah NG & Khatib AM (2008) Selective inhibition of proprotein convertases represses the metastatic potential of human colorectal tumor cells. Journal of Clinical Investigation 118: 352–363

Shigeta K, Datta M, Hato T, Kitahara S, Chen IX, Matsui A, Kikuchi H, Mamessier E, Aoki S, Ramjiawan RR, et al (2020) Dual PD-1 and VEGFR-2 blockade promotes vascular normalization and enhances anti-tumor immune responses in HCC HHS Public Access. Hepatology 71: 1247–1261

Singh K, Foster MW, Violette MJ, Corcoran AM, Hotchkiss KM, Railton CO, Blandford EE, Blethen KE, Thomas EL, McIntosh WC, et al (2025) A surgical window of opportunity trial evaluating the effect of the PCSK9 inhibitor evolocumab on tumoral MHC-I expression and CD8+ infiltration in glioma. Scientific Reports 2025 15:1 15: 37112-

Sun X, Essalmani R, Day R, Khatib AM, Seidah NG & Prat A (2012) Proprotein Convertase Subtilisin/Kexin Type 9 Deficiency Reduces Melanoma Metastasis in Liver. Neoplasia 14: 1122–IN5

Sun Y, Zhou P, Qian J, Zeng Q, Wei G, Li Y, Liu Y, Lai Y, Zhan Y, Wu D, et al (2025) Spermine synthase engages in macrophages M2 polarization to sabotage antitumor immunity in hepatocellular carcinoma. Cell Death Differ 32: 573–586

Tian P, Wu Q, He D, Zhao W, Luo L, Jia Z, Luo W, Lv X, Liu Y, Wang Y, et al (2025) Breast cancer induces CD62L+ Kupffer cells via DMBT1 to promote neutrophil extracellular trap formation and liver metastasis. Cell Discov 11

Vidal-Vanaclocha F (2008) The Prometastatic Microenvironment of the Liver. Cancer Microenvironment 1: 113

Vidal-Vanaclocha F. (2011) Architectural and functional aspects of the liver with implications for cancer metastasis. Brodt P. (ed) Netherlands: Springer

Wang L, Li S, Luo H, Lu Q & Yu S (2022) PCSK9 promotes the progression and metastasis of colon cancer cells through regulation of EMT and PI3K/AKT signaling in tumor cells and phenotypic polarization of macrophages. Journal of Experimental and Clinical Cancer Research 41: 1–21

Wong CC, Wu JL, Ji F, Kang W, Bian X, Chen H, Chan LS, Luk STY, Tong S, Xu J, et al (2022) The cholesterol uptake regulator PCSK9 promotes and is a therapeutic target in APC/KRAS-mutant colorectal cancer. Nature Communications 2022 13:1 13: 1–15

Xiao Z, Chen D, Mulder F, Song S, van der Wouden PE, Cool RH, Melgert BN, Poelarends GJ & Dekker FJ (2022) 4-Iodopyrimidine Labeling Reveals Nuclear Translocation and Nuclease Activity for Both MIF and MIF2. Chemistry 28

Ye P, Xi Y, Huang Z & Xu P (2020) Linking Obesity with Colorectal Cancer: Epidemiology and Mechanistic Insights. Cancers (Basel) 12

Zheng M, Meng H, Li Y, Shi J, Han Y, Zhao C, Chen J, Han J, Liang J, Chen Y, et al (2023) S100A11 Promotes Metastasis via AKT and ERK Signaling Pathways and Has a Diagnostic Role in Hepatocellular Carcinoma. Int J Med Sci 20: 318–328

