## Supplementary figure 1 and 2 and Supplementary Table 1 for "PCSK9 Exhibits Novel Nuclear Localization in LSEC and Its Targeting with Bioinspired Nanoparticles Reduces Colorectal Liver Metastasis"

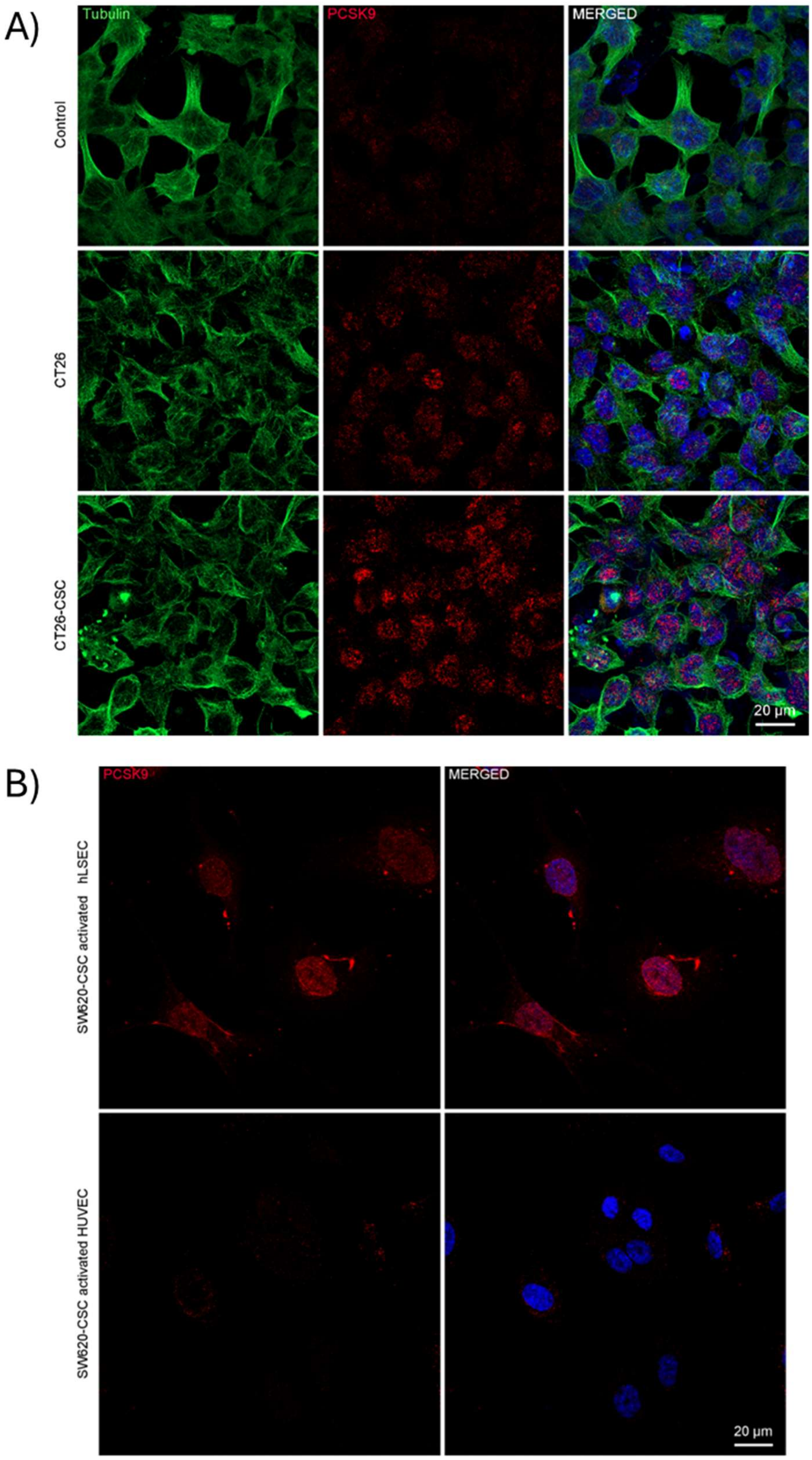


**Supplementary Figure 1. Analysis of the subcellular localization of PCSK9 in murine LSECs and HUVECs**. A) Immunofluorescence staining of PCSK9 and tubulin in murine LSEC cultures under basal conditions and following activation with conditioned media from differentiated metastatic colorectal tumor cells and colorectal cancer stem cells (CSCs). B) Immunofluorescence staining of PCSK9 in human LSECs and human umbilical vein endothelial cells (HUVECs).

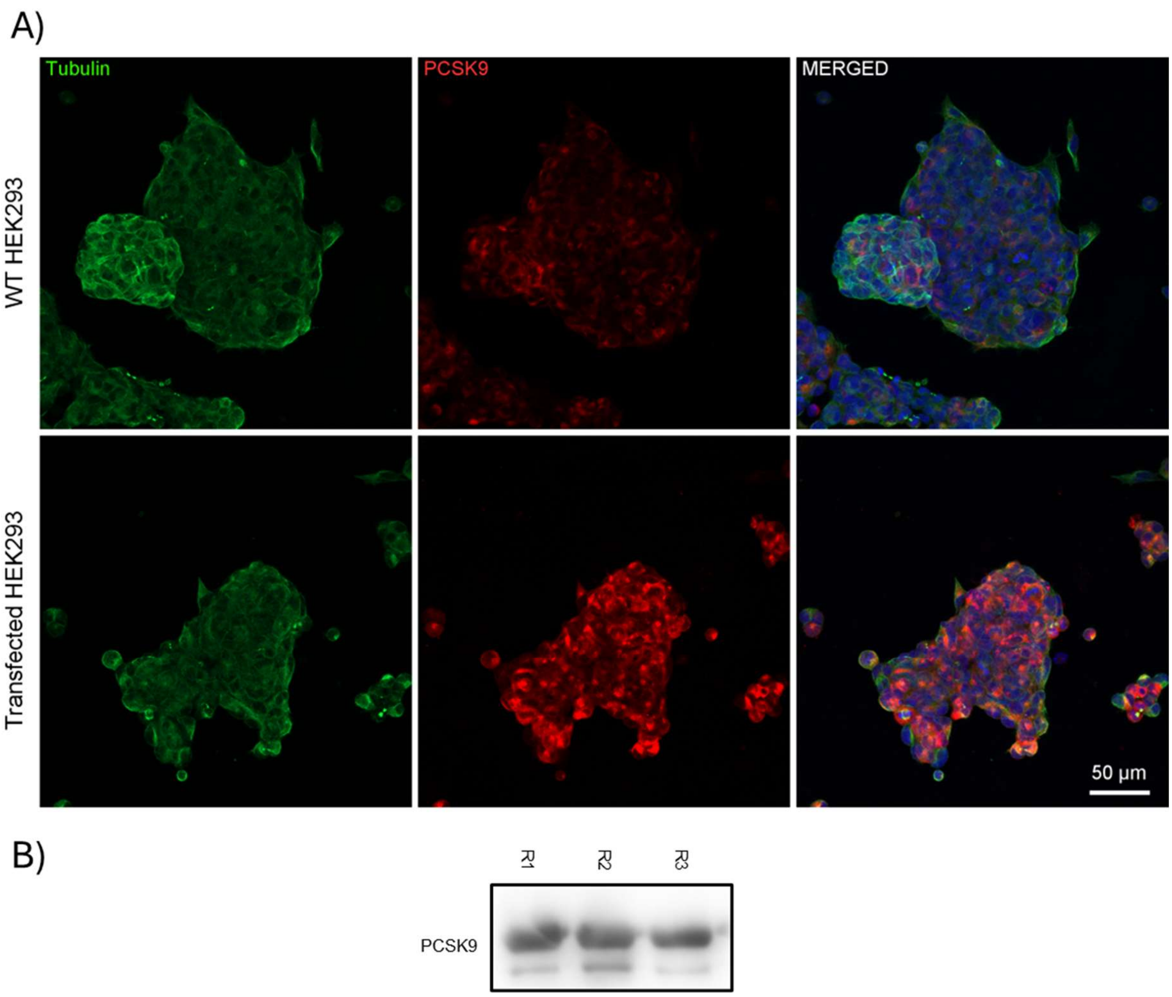


**Supplementary Figure 2. Validation of PCSK9 transfection and extract quality.** A) Immunofluorescence staining of tubulin and PCSK9 in wild-type HEK293 cells and HEK293 cells transfected with the PCSK9 plasmid (pCMV-PCSK9-FLAG). B) Western blot analysis of the three replicate co-immunoprecipitation (Co-IP) extracts that were subsequently analyzed by mass spectrometry

**Supplementary Table 1.** Dynamic light scattering (DLS) characterization of nanoparticles administered in vivo, including Z-average diameter and polydispersity index (PDI) for each injection day.

| Injection | Sample | Dilution | Z Average (nm) | PDI |
|---|---|---|---|---|
| 1st treatment injection | Empty NP | 50 fold in H2O | 142,30 ± 0,49 | 0,124 ± 0,020 |
| | | 50 fold in 5% Glucose | 248,63 ± 9,50 | 0,089 ± 0,040 |
| | siCtl NP | 50 fold in H2O | 146,10 ± 1,70 | 0,106 ± 0,004 |
| | | 50 fold in 5% Glucose | 609,30 ± 38,48 | 0,250 ± 0,015 |
| | siPCSK9 NP | 50 fold in H2O | 159,03 ± 1,60 | 0,104 ± 0,005 |
| | | 50 fold in 5% Glucose | 206,83 ± 2,11 | 0,121 ± 0,013 |
| 2nd treatment injection | Empty NP | 50 fold in H2O | 151,70 ± 5,69 | 0,109 ± 0,020 |
| | | 50 fold in 5% Glucose | 163,83 ± 4,80 | 0,056 ± 0,040 |
| | siCtl NP | 50 fold in H2O | 129,13 ± 0,40 | 0,138 ± 0,010 |
| | | 50 fold in 5% Glucose | 145,40 ± 1,50 | 0,110 ± 0,010 |
| | siPCSK9 NP | 50 fold in H2O | 155,43 ± 2,06 | 0,131 ± 0,030 |
| | | 50 fold in 5% Glucose | 204,10± 2,28 | 0,236 ± 0,009 |
| 3rd treatment injection | Empty NP | 50 fold in H2O | 157,60 ± 0,98 | 0,099 ± 0,010 |
| | | 50 fold in 5% Glucose | 181,50 ± 4,15 | 0,108 ± 0,013 |
| | siCtl NP | 50 fold in H2O | 149,90 ± 1,80 | 0,130 ± 0,002 |
| | | 50 fold in 5% Glucose | 167,83 ± 1,47 | 0,074 ± 0,020 |
| | siPCSK9 NP | 50 fold in H2O | 163,30 ± 0,90 | 0,180 ± 0,028 |
| | | 50 fold in 5% Glucose | 173,93 ± 4,75 | 0,094 ± 0,016 |
| 4th treatment injection | Empty NP | 50 fold in H2O | 143,26 ± 1,00 | 0,128 ± 0,004 |
| | | 50 fold in 5% Glucose | 204,10 ± 7,07 | 0,223 ± 0,006 |
| | siCtl NP | 50 fold in H2O | 169,10 ± 0,36 | 0,110 ± 0,005 |
| | | 50 fold in 5% Glucose | 171,80 ± 3,82 | 0,250 ± 0,007 |
| | siPCSK9 NP | 50 fold in H2O | 142,73 ± 0,98 | 0,152 ± 0,020 |
| | | 50 fold in 5% Glucose | 186,23 ± 9,28 | 0,128 ± 0,020 |
